# Sex disparities in influenza: a multiscale network analysis

**DOI:** 10.1101/2021.03.25.437108

**Authors:** Chang Wang, Lauren P. Lashua, Chalise E. Carter, Scott K. Johnson, Minghui Wang, Ted M. Ross, Elodie Ghedin, Bin Zhang, Christian V. Forst

## Abstract

Sex differences in the pathogenesis of infectious diseases due to differential immune responses between females and males have been well documented for multiple pathogens. However, the molecular mechanism underlying the observed sex differences in influenza virus infection remains poorly understood. In this study, we used a network-based approach to characterize the blood transcriptome collected over the course of infection with influenza A virus from female and male ferrets to dissect sex-biased gene expression. We identified significant differences in the temporal dynamics and regulation of immune responses between females and males. Our results elucidate sex-differentiated pathways involved in the unfolded protein response (UPR), lipid metabolism, and inflammatory responses, including a female-biased IRE1/XBP1 activation and male-biased crosstalk between metabolic reprogramming and IL-1 and AP-1 pathways. Overall, our study provides molecular insights into sex differences in transcriptional regulation of immune responses and contributes to a better understanding of sex bias in influenza pathogenesis.

## INTRODUCTION

Sex-related differences shaped by multidimensional biological characteristics that define females and males exert considerable influence on the pathogenesis of various diseases [1], including autoimmune diseases [2], cancers [3–5], and infectious diseases caused by diverse pathogens [6–8]. Each infectious disease exhibits a distinct pattern of sex bias in the prevalence, intensity, and outcome of infections [6, 9], as well as in the responses to antiviral drugs and vaccines [10, 11]. For example, females have a higher fatality rate following exposure to influenza A viruses (IAV) [12] and more robust antibody responses and adverse reactions after vaccination than males [13–15], while accumulating evidence suggests a male bias in COVID-19 mortality [16, 17]. Thus an in-depth understanding of sex differences in the pathogenesis of diseases and consideration in the rational design of prophylactic and therapeutic strategies represent an important step towards personalized medicine.

The observed sex differences in disease pathogenesis have been primarily attributed to the differences in the innate and adaptive immune responses between females and males. In both arms of immunity, the sexes differ in multiple aspects, including the detection of pathogen nucleic acids by pattern recognition receptors (PRRs) such as the Toll-like receptors (TLRs), the number and activity of immune cells, and the production of cytokines and chemokines (reviewed in [1, 18]). These intricate differences in immune functions between sexes have a powerful impact on infectious disease pathogenesis. For example, an augmented response to pathogens in females allows better control and clearance of pathogens while promoting increased immunopathology [6, 7, 10]. During influenza infections, female mice exhibited a more robust induction of pro-inflammatory cytokines and chemokines in their lungs, including TNF-*α*, IFN-*β*, IL-6, and CCL2, accompanied by greater weight loss, hypothermia, and mortality than male mice [19].

The etiology underlying sex differences in immunity involves intrinsic and extrinsic factors that exert a combinatorial effect on the immune system’s functioning. Although the microbiome [20–23] and nutritional status [24–29] have been implicated in modulating immune responses, sex hormones and genetic mediators are the most widely appreciated factors shaping differential immunity between females and males. In addition to the profound effects of sex hormones that have been extensively demonstrated, genetic differences attributed to immune-related genes and microRNAs (miRNAs) that are located on the sex chromosomes also play an important role in determining the distinct immune responses between sexes, especially in prepubertal children, postmenopausal females and age-matched males (reviewed in [1, 6, 7, 18, 30–33]). However, the pathways and cellular responses that mediate the differences in response to influenza infection have not been fully elucidated. Moreover, the confounding effects of hormone and genetic factors in intact animals and human populations impose a challenge in resolving the mechanism of sex differences in influenza pathogenesis.

To address the molecular mechanisms underlying sex differences in influenza infection responses, we assessed the transcriptomic dynamics of blood cell responses in neutered female and male ferrets throughout the infection. We detected a temporal shift in mounting immune responses against viral infection between females and males. Using a multiscale network-based approach, we further identified pathways and cellular processes commonly induced in both sexes and those uniquely regulated in females or males. Our data revealed gene regulatory pathways that were differentially regulated between sexes and likely involved in marked sexual differences in influenza virus-induced pathogenesis, providing molecular and functional insights for functional evaluation and development of therapeutic strategies.

## RESULTS

### An integrated network approach for systematic characterization of the immune response to influenza infection in females and males

To evaluate the genetic factors mediated sex differences in the immune response to influenza virus infection, we conducted a study with neutered female and male ferrets. We examined the transcriptional response in their blood cells over the course of the infection (**Fig. 1a**). Briefly, whole blood samples were collected from spayed adult female and castrated adult male ferrets infected with the influenza virus A/CA/07/2009 (H1N1pdm09) strain over a period of 1 to 8 (for males) or 14 (for females) days post-infection (dpi). Samples from infected animals and uninfected control samples, including those collected before the infection from infected animals or prior to and post-infection from uninfected individuals, were used to serve as the baseline for the comparison of infection responses within each sex. The whole blood transcriptome was profiled to assess the impact of genetic sex on immune response regulation without the additive influence of sex hormones. Given the time frame of sample collection, this study primarily focused on identifying sex differences in innate immunity and the early phase of adaptive immunity [34].

**Figure 1.**
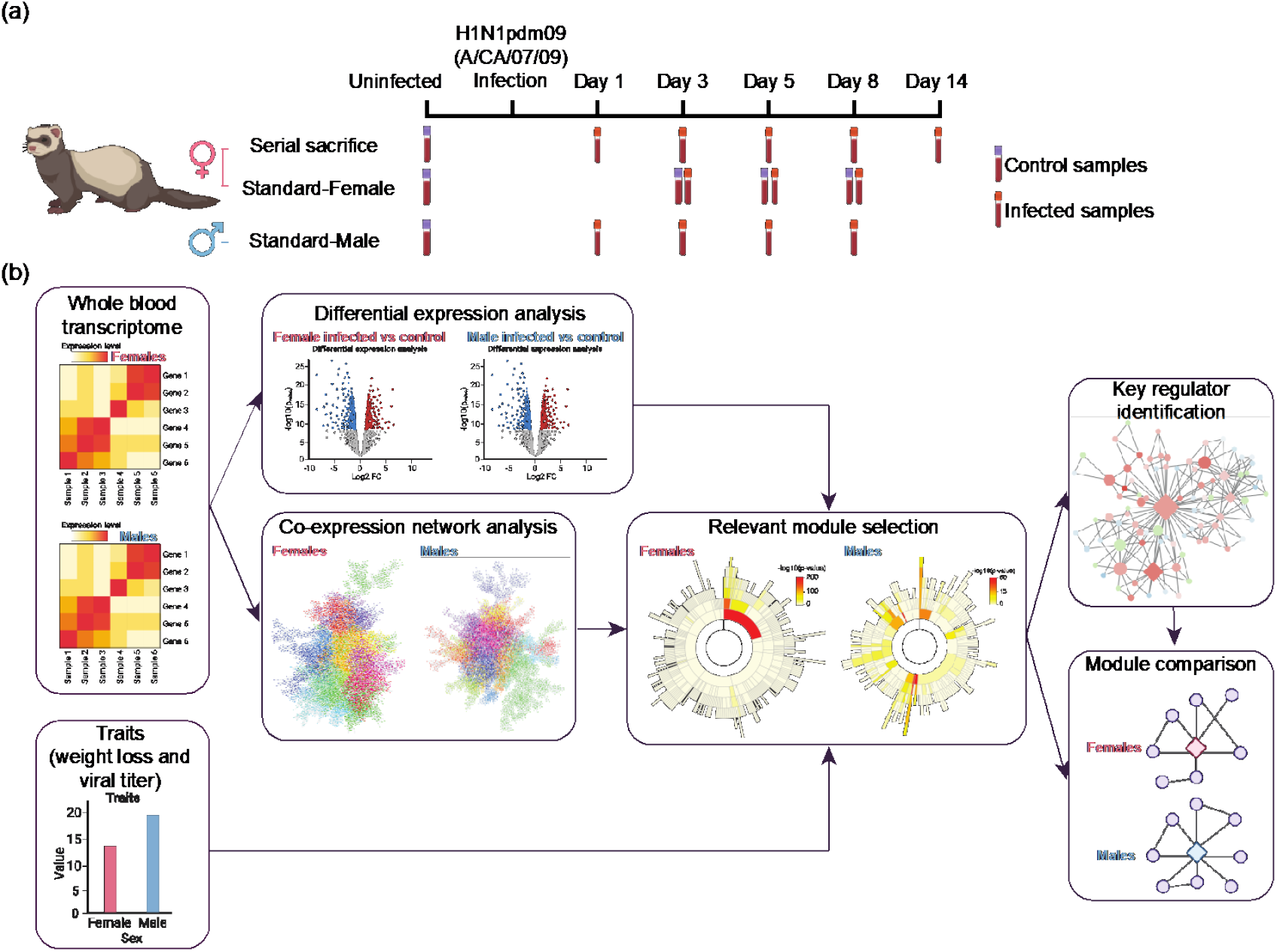
Overview of study design and analysis scheme. (**a**) Sample collection strategy of adult female and male ferrets infected with influenza A/CA/07/09 (H1N1pdm09) virus. In the serial sacrifice study of female ferrets (n = 45), blood samples were collected at the uninfected baseline (n = 26) and days 1 (n = 10), 3 (n = 12), 5 (n = 8), 8 (n = 7), and 14 (n = 8) post-infection from animals sacrificed at a given day. For the standard study of female ferrets (n = 9), samples were collected at baseline and from naïve (n = 4) and infected (n = 5) animals at days 3, 5, and 8 post-infection. Similarly, for the standard study of male ferrets (n = 34), samples were collected at baseline (n = 33) and at days 1, 3, 5, and 8 post-infection (n = 30). (**b**) Framework for systematic characterization of sex difference in the immune response to influenza infection with whole blood transcriptome. Transcriptomic data generated from infected and control samples of each sex were used for differential expression analysis and Multiscale Embedded Gene Co-expression Network Analysis (MEGENA) separately. Disease-relevant modules of co-expressed genes were then identified by the enrichment of differentially expressed genes (DEGs) and the association with physiological traits. Key regulators in those selected modules were predicted using the adopted Fisher’s inverse Chi-square approach in MEGENA (see **Methods**). Pairwise comparisons further determined similarities of module composition, key regulators, and within-module correlation pattern between female and male modules (see **Methods**).

To systematically characterize sex differences in the immune response to influenza virus infection, we used a co-expression network-based approach that integrated transcriptional patterns with physiological traits and enabled the identification of sex-specific gene expression signatures (**Fig. 1b**). Specifically, differential expression analysis was carried out within each sex by comparing the infected samples with the uninfected controls at each time point post-infection. A Multiscale Embedded Gene Co-expression Network Analysis (MEGENA) was also performed [35] with the transcriptomic data for each sex separately and in parallel. We further prioritized MEGENA network modules based on their enrichment for differentially expressed genes (DEGs). We assessed their functions with the Molecular Signature Database (MSigDB) and blood cell-type-specific gene sets. Disease-relevant modules were then correlated with pathological characteristics, including weight loss and viral load (**Fig. S1**), and selected for comparison of response signatures and co-expression patterns between sexes. Key regulators determined by multiscale hub analysis (MHA, see *Methods*), as well as module compositions and within-module correlation patterns in selected modules, were further investigated to elucidate shared or sex-unique processes and regulatory patterns.

### Differential expression analysis reveals global temporal differences in immune response dynamics between sexes

Following the recognition of viruses, various immune cells equipped with versatile strategies to mount a potent antiviral response are mobilized to establish physical and chemical barriers against viral infection. To determine the global dynamics of blood cell response in females and males over time, we first carried out differential expression analyses with the whole blood transcriptome data collected over 8- or 14-day periods post influenza infection by comparing with sex-matched uninfected controls. Among different sets of DEGs identified over time in females and males (**Tables S1** and **S2**), the most significant overlaps between any two given sets were observed at 3 and 5 dpi within or between sexes, followed by those detected between 3-5 dpi in each sex and 1 dpi in females or 8 dpi in males (**Fig. 2a** and **Fig. S2**). It is noteworthy that DEGs at 14 dpi in females did not share significant overlaps with any other sets (**Fig. 2a**), which suggests a largely subsided initial immune response by 14 dpi. This finding is consistent with the reported kinetics of an early innate phase in biphasic immune responses during influenza infections in ferrets [34]. To further understand the biological processes associated with DEGs during viral infection, we explored the enrichment of MSigDB gene sets with up- and down-regulated DEGs at each time point over 8 days in each sex. As expected, there was an enrichment of genes involved in various aspects of the immune response to viral infection in both sexes, including the innate immune response and cytokine signaling — particularly type I (*α/β*) and II (*γ*) interferon (IFN) signaling — hemostasis, the urokinase-type plasminogen activator (uPA) and its receptor (uPAR) mediated signaling, DNA replication and the cell cycle (**Fig. 2b**). These processes exhibited differential temporal kinetics of activation, with IFN signaling being initiated immediately upon infection. However, cell cycle-related processes were revealed later in the infection (**Fig. 2b**). We also detected enrichment of down-regulated genes during infection in the signatures of T-cell functioning and IL-17 pathways in both sexes and several generic cellular processes related to transcription and translation in males (**Fig. 2b** and **Tables S3** and **S4**).

**Figure 2.**
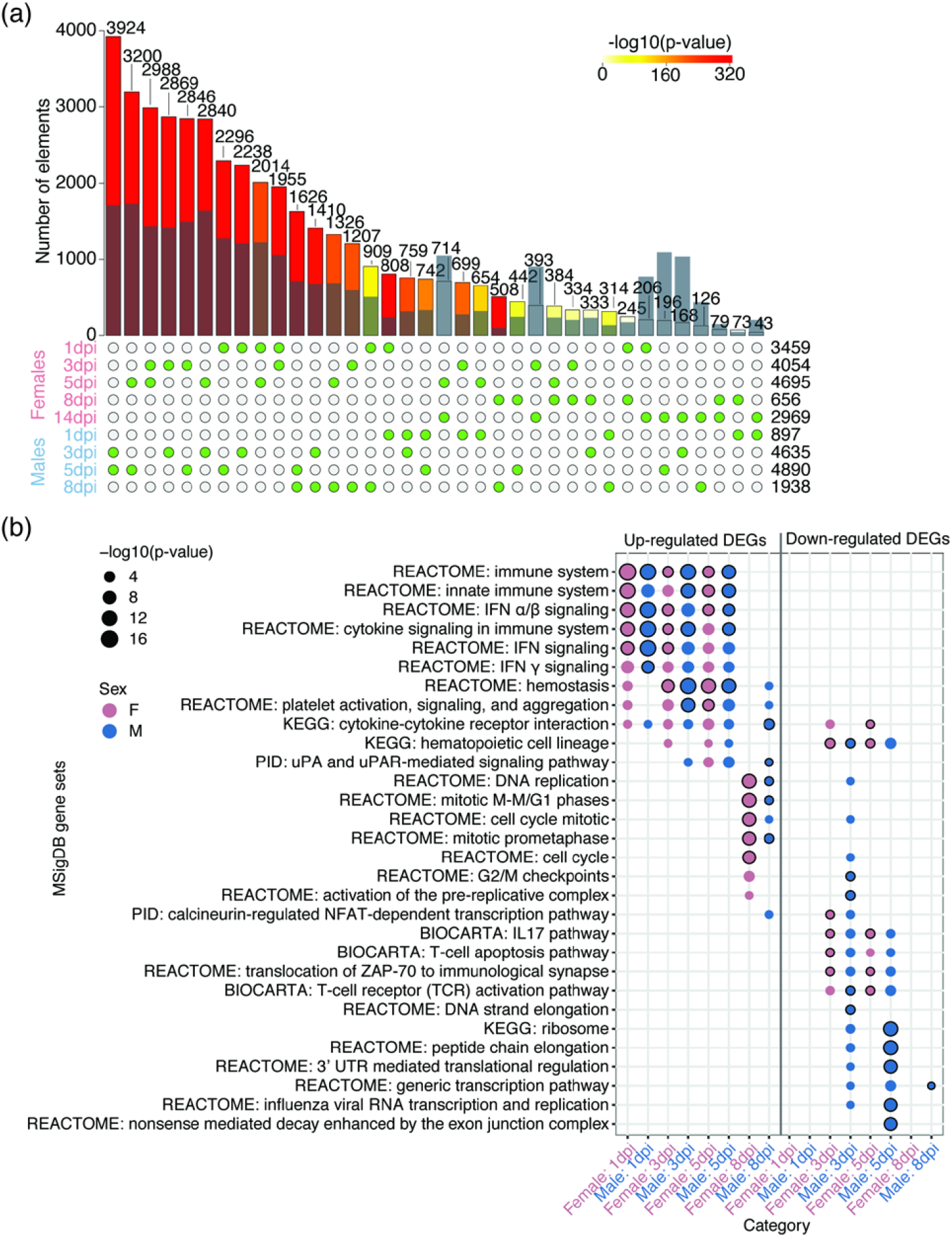
Global dynamics of blood cell response following infection in both sexes. (**a**) Pairwise comparison of differentially expressed genes (DEGs) at each time point in both sexes. DEGs were obtained by comparing the infected samples with their controls for each sex. The solid bars are colored by -log10(p-value) generated with the super exact test and the numbers at the top show the observed intersects of DEGs between a pair of DEG sets. The semi-transparent blue bars overlaying the solid bars indicate the expected intersects. The green dots below denote two sets of DEGs used in the comparisons. The numbers to the right of the dot matrix indicate the number of DEGs in a given set. Detailed comparisons of up- and down-regulated DEGs between sexes over time can be found in **Fig. S2**. DEG lists can be found in **Tables S1** and **S2**. (**b**) Molecular Signatures Database (MSigDB) gene sets associated significant DEGs at each time point in both sexes. The size of dots indicates adjusted p-values for gene set enrichment tests, and the color denotes the sex. The top 5 gene set signatures enriched in each DEG set were shown and highlighted with strokes around the dots. Detailed information can be found in **Tables S3** and **S4**.

To further investigate the temporal dynamics of infection-responsive processes in both sexes, we examined the fold change (FC) of DEGs associated with the enriched MSigDB gene sets over time. We found more robust activation of type I and II IFN signaling pathways in females than males, as many genes were immediately and more strongly activated at 1 dpi in females (**Fig. 3a–3b**). A temporal shift in the expression kinetics between sexes was also evident in platelet activation, signaling, and aggregation, which were important in maintaining hemostasis and modulating inflammation [36]. Many genes involved in platelet activity were immediately activated at 1 dpi and substantially subsided by 8 dpi in females, in contrast to an attenuated activation at 1 dpi and continuous expression beyond 8 dpi in males (**Fig. 3c**), further suggesting a more rapid response in females than males. Interestingly, we also detected a DEG, *GLIPR1L2* (GLI pathogenesis-related 1 like 2), with extreme temporal shift as manifested by inverse alteration kinetics between the sexes (r = −0.9993, p = 0.0238). *GLIPR1L2* was down-regulated at 1-3 dpi and returned to baseline expression by 5 dpi in the females but was continuously down-regulated from 3 dpi onwards in males (**Fig. 3d**). Although the function of *GLIPR1L2* in viral infections remains unclear, it has been shown to reside in the same gene cluster with *GLIPR1* (GLI pathogenesis-related protein 1) and *GLIPR1L1* and be targeted by the tumor suppressor p53 during cancer development and progression [37]. Taken together, these results indicate temporal differences in the immune response dynamics between the sexes, marked by a prompt response in females compared with a lagged response in males.

**Figure 3.**
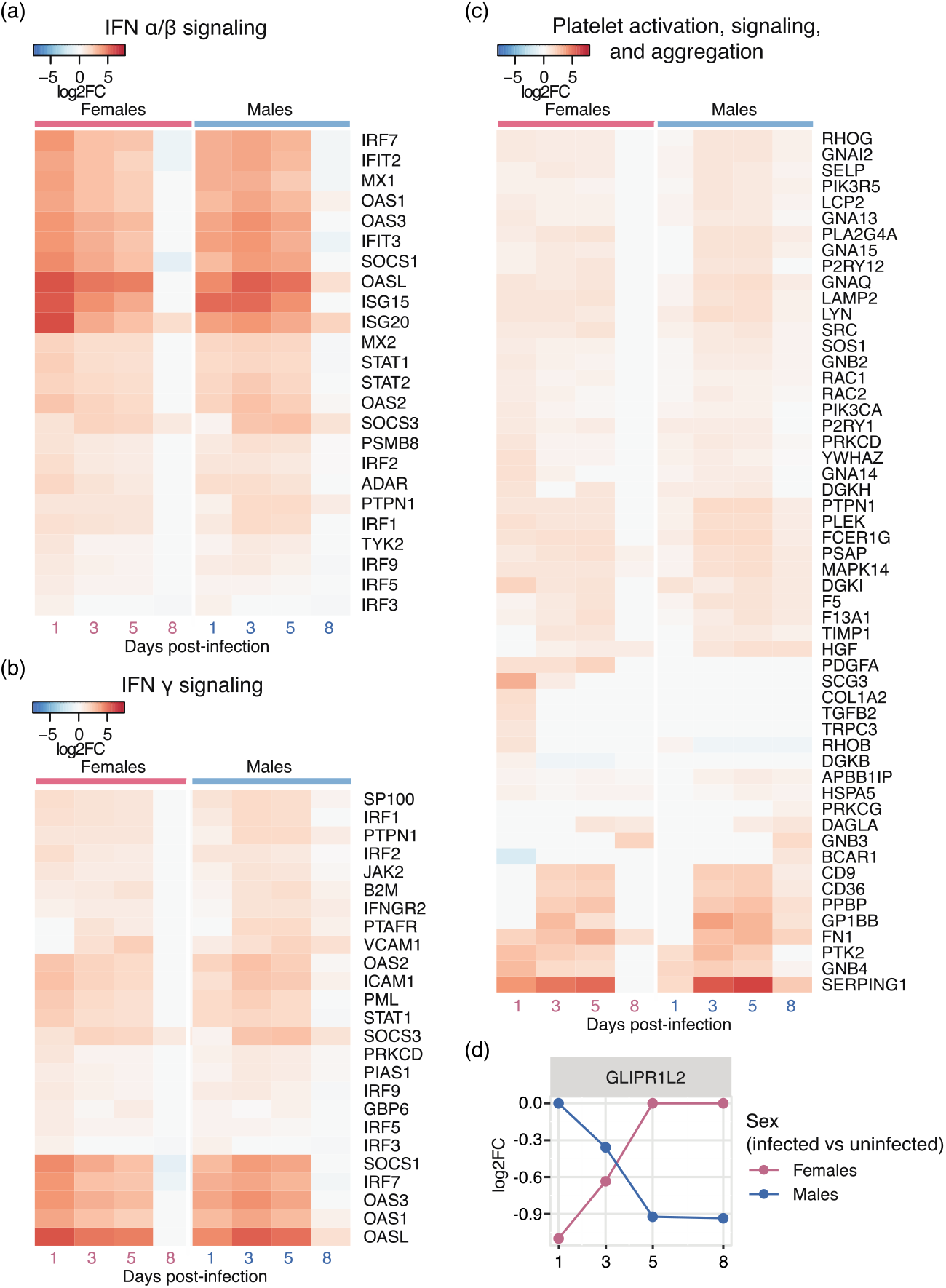
Temporal differences in dynamics of response to infection between sexes. (**a-d**) log2(fold change (FC)) of differentially expressed genes (DEGs) involved in (**a**) interferon (IFN) *α/β* signaling, (**b**) IFN *γ* signaling, and (**c**) platelet activation, signaling, and aggregation over the course of the infection in both sexes. DEGs that were considered significant (i.e., with a log2(FC) ≥ log2(1.5) or ≤ -log2(1.5)) at one or more time points in females or males are shown. (**d**) log2(FC) of *GLIPR1L2* over time. The colors denote different comparisons within each sex.

### Co-expression network analysis identifies shared and sex-unique expression patterns in the immune response

Although differential expression analysis enabled a global overview of the immune response in both sexes over the course of the infection, it did not provide the resolution for identifying gene subsets with sex-unique expression patterns. To dissect the sex-biased factors, we next sought to employ a network-based approach in search of consistent and unique gene expression patterns between sexes with increased granularity. With MEGENA, we constructed multiscale gene co-expression networks for females and males separately (**Tables S5** and **S6**) and prioritized modules of co-expressed genes in each sex based on their enrichment of DEGs and coherent biological processes and pathways. To assess the direct link between modules and pathogenesis, we also established a correlation between modules and physiological features, including viral titers in the nasal washes and weight change (**Fig. S1**). Overall, we identified modules enriched for up- or down-regulated DEGs, or both, at one or more time points in each sex (**Fig. 4a–4b** and **Fig. S3** and **Tables S7** and **S8**). Those modules were enriched in a diverse array of gene set signatures involved in distinct but related aspects of IFN signaling, inflammation, immune cell migration, hemostasis, and apoptosis induction and clearance exerted by various blood cells (**Fig. 4c**), revealing more specific processes than those associated with DEGs globally. Some modules were also positively associated either with viral load in one or both sexes or with weight loss in males (**Fig. 4c** and **Fig. S4**). For example, viral load-related modules in both females and males were consistently enriched in gene set signatures of IFN signaling, primarily type I IFN signaling. In contrast, viral load-related modules in females alone were associated with specific inflammation-related pathways, lysosome-related processes, the unfolded protein response (UPR), and hemostasis. Those only in males were associated with apoptosis-related pathways (**Fig. 4c** and **Fig. S4**). Although we did not detect any weight loss-related modules in females, those in males exhibited signatures of various generic processes and pathways, including transcription, translation and its regulation, the AHSP pathway, nuclear *β*-catenin pathway, 4-1BB dependent immune response pathway, stathmin pathway, the primary immunodeficiency pathway, porphyrin metabolism, and generation of second messengers (**Fig. 4c** and **Fig. S4**), suggesting a complex relationship between weight loss and underlying molecular processes. Moreover, enrichment of blood cell-type-specific signatures was also identified in several modules in each sex, including (i) the T-cell signature related to either the translocation of ZAP-70 to the immunological synapse pathway in female modules M35, M196, and M524 and in the male module M127, or to (ii) the generation of second messenger molecules pathway in the male module M15, and (iii) the CD19^+^ B-cell signature related to primary immunodeficiency pathway in the male modules M17, M134, M356, and M523 (**Tables S7** and **S8**). These results elucidate a comprehensive landscape of complex immune responses elicited by diverse types of immune cells. Although both sexes have a common arsenal of strategies as immune system response, females and males further employ different pathways and processes contributing to the fine granularity of sex disparity in such a host-pathogen system.

**Figure 4.**
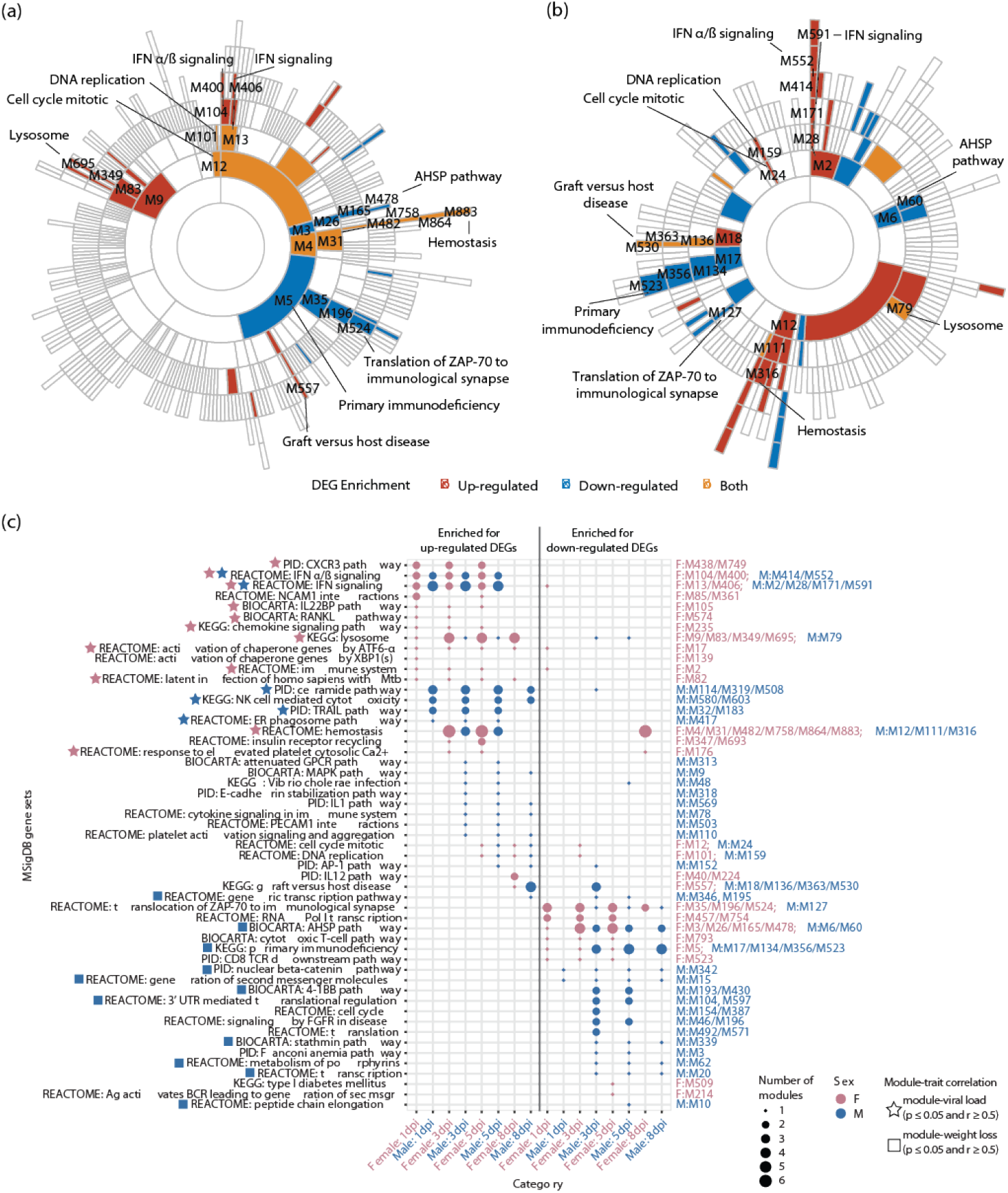
Co-expressed gene modules enriched in infection-responsive genes in both sexes. (**a-b**) Sunburst plots showing the hierarchy of all the modules and those enriched in differentially expressed genes (DEGs) in the (**a**) female and (**b**) male co-expression networks, respectively. Modules are colored by the enrichment in up- (red) or down- (blue) regulated DEGs or both (yellow). Modules enriched in the Molecular Signatures Database (MSigDB) gene sets observed in both sexes are highlighted. Detailed information about each module’s composition can be found in **Tables S5** and **S6,** and information about modules enriched in DEGs at each time point in **Fig. S3** and **Tables S7** and **S8**. (**c**) Bubble plot showing the MSigDB gene sets associated with modules enriched in DEGs in both sexes. The size of the bubbles indicates the number of modules enriched in an MSigDB gene set, and the color denotes the sex. The modules and their hierarchy were shown on the right side of the plot and colored by sex. The asterisks and squares next to the MSigDB gene sets indicate the presence of correlation between at least one module among those associated with the corresponding MSigDB gene set and physiological traits (i.e., asterisks for viral load and squares for weight loss). Only the correlations with p-value ≤ 0.05 and coefficient ≥ 0.5 were included. Detailed information can be found in **Fig. S4**.

To identify the molecular factors mediating sex differences in the immune response to infection, we performed an in-depth comparison of the structural and functional relationships between modules across the female and male networks. Given that some processes and pathways were consistently detected over the 8-day period in the female and male networks, we first asked if the modules enriched in identical gene set signatures in both sexes shared similar members and key regulators. These shared signatures depicting temporal dynamics of the immune response included (i) IFN signaling (primarily type I IFN), lysosome-related processes, hemostasis, mitotic cell cycle, and DNA replication, and graft versus host disease signature, which exhibited gradual activations over time, and (ii) primary immunodeficiency signature, AHSP pathway, and translocation of ZAP-70 to the Immunological synapse, which were generally suppressed in both sexes (**Fig. 4**), similar to those observed in DEGs. We conducted pairwise comparisons between each module in the female and male networks with Fisher’s Exact Test (FET) to determine the similarity of modules and their key regulators (**Table S9**). We identified significant overlaps between the modules associated with those shared signatures in both sexes (**Table S9**). However, the key regulators in modules with matched signatures shared limited overlap (**Fig. S5** and **Table S9**). These results suggest similar transcriptional programs in both sexes upon infection are coordinated by different regulators in females and males.

To further pinpoint the sex-specific gene co-expression patterns that implicated distinct functional signatures, we next focused on the modules associated with sex-unique gene set signatures and determined the uniqueness of their module memberships in both sexes. We did a module member similarity test (i.e., FET) along with a differential gene correlation analysis (DGCA) [38] with genes in modules enriched in processes or pathways uniquely observed in one sex. DGCA assesses if those genes possessed a different regulatory relationship in their counterpart module initially determined by FET in the other sex (**Table S10**). We detected one female module, M139, which had a female-unique composition determined by FET (**Table S9**), and that was associated with the activation of chaperone genes by XBP1(s) (**Fig. 4c** and **Table S7**). We also identified two male modules, M152 and M569, which had male-unique regulatory relationships identified with DGCA between module counterparts (**Table S10**) and were associated with the AP-1 (activator protein-1) and IL-1 pathways, respectively (**Fig. 4c** and **Table S8**). The female-specific module M139 contained *XBP1* (X-box binding protein 1) as its key regulator (**Fig. 5a**). The majority of the genes within this module exhibited a rapid and strong induction immediately following infection at 1 dpi in females (**Fig. S6a**) compared with a milder induction in males (**Fig. 5b**), suggesting a prompt and potent activation of the IRE1/XBP1 branch of the UPR [39, 40] during infections, specifically in females. The male-specific module M152 was enriched in later-activated DEGs, including two key regulators *HPGD* (15-Hydroxyprostaglandin dehydrogenase) and *MS4A2* (the *β* chain of the high-affinity IgE receptor (Fc*ε*RI*β*)), which were down-regulated at 3 dpi and up-regulated at 5-8 dpi in both sexes (**Fig. 5c–5d** and **Fig. S6b**) and likely linked to Fc*ε* RI- and prostaglandin E_2_ (PGE_2_)-mediated inflammatory responses [41–45]. The other male-specific module, M569, enriched in strongly up-regulated DEGs at 3-8 dpi in males (**Fig. S6c**) compared with that in females, contained two key regulators *ACSL3* (acyl-CoA synthetase long-chain family member 3) and *IL18RAP* (IL-18 receptor accessory protein) (**Fig. 5e–5f**), implying a link between lipid metabolism and inflammation [46–51]. Taken together, these results revealed sex-dependent regulatory relationships and the extent of the modulation of genes involved in IRE1-XBP1, AP-1, and IL-1 pathways upon infection, suggesting sex differences in specific aspects of the UPR, lipid metabolism, and inflammation independent of sex hormones.

**Figure 5.**
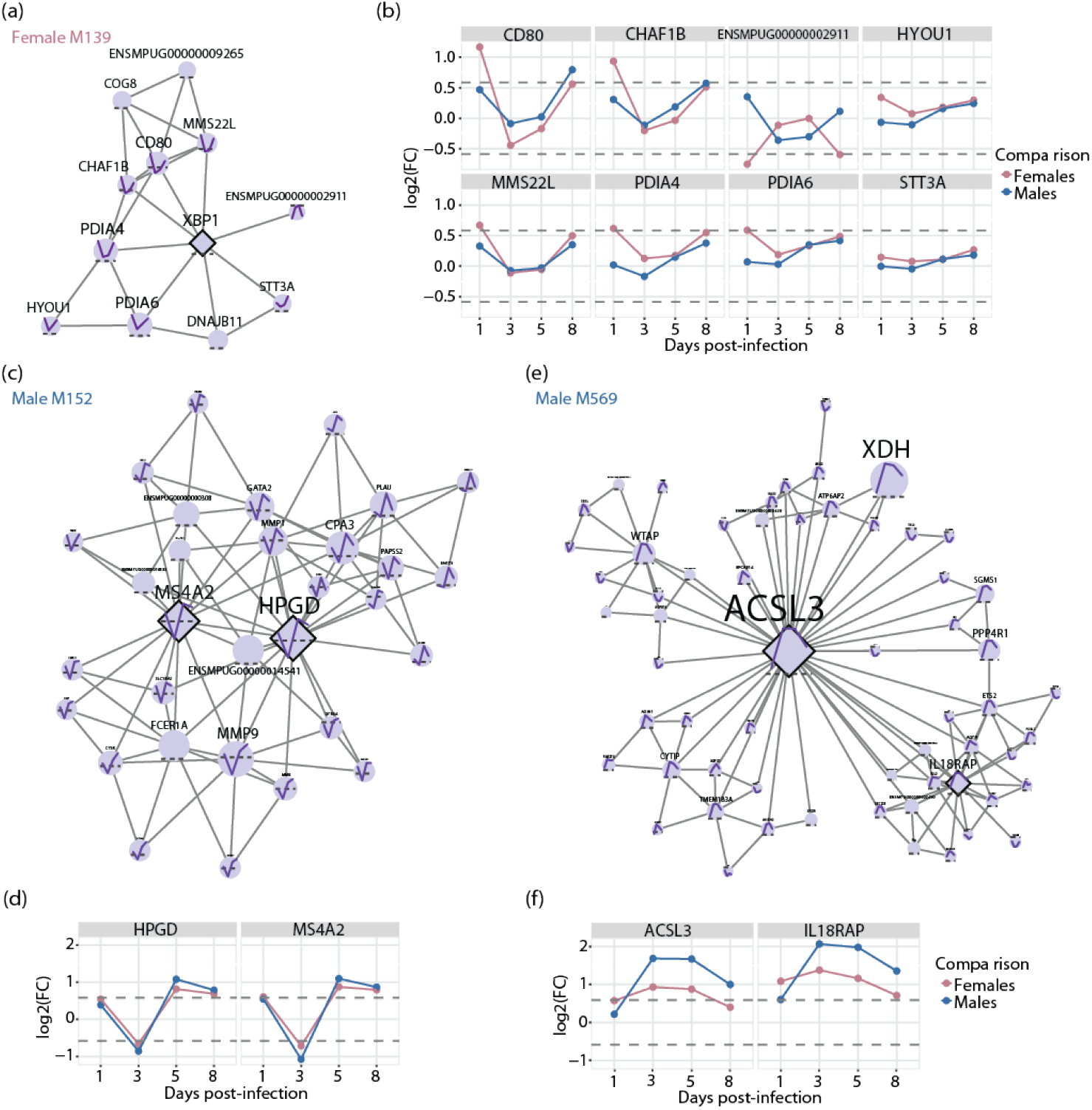
Networks of sex-unique modules and the temporal expression pattern of their key regulators and members. (**a**, **c**, and **e**) Networks of co-expressed genes in (**a**) a female-specific module M139 and (**c** and **e**) two male-specific modules M152 and M569. Node shapes denote the module membership (diamonds for key regulators and circles for regular members), and their sizes are proportional to the node strength. The solid purple lines in the center of nodes show the log2(fold change (FC)) of a given differentially expressed gene (DEG), and the dashed grey lines indicate zero. (**b**, **d**, and **f**) log2(FC) of (**b**) DEGs in the female-specific module M139, or that of the key regulators in the male-specific modules (**d**) M152 and (**f**) M569. The colors indicate the comparison within each sex (i.e., pink for females and blue for males). Detailed information about DEGs in each module over time can be found in **Fig. S6**.

## DISCUSSION

A comprehensive understanding of the molecular mechanism underlying sex differences in response to viral infections has been either largely limited to sex chromosome-linked genes or hindered by the presence of confounding factors, especially in human populations. In this study, we systematically characterized sex differences in the transcriptional landscape of blood cell responses to influenza virus infection in neutered ferrets, precluding the impact of other factors. We detected temporal and regulatory differences in the processes commonly induced in both sexes, as evidenced by a more rapid response in females than males and different key regulators orchestrating those shared responses. With a network-based approach, we further identified sex-specific modules of co-expressed genes that exhibited unique regulatory relationships in each sex. Those sex-unique modules revealed a rapid UPR activation via the IRE1/XBP1 pathway upon infection in a female-biased manner and a male-specific regulation of inflammatory responses mediated by a cascade involving Fc*ε*RI, AP-1, and PGE_2_, and the IL-1 family signaling associated with lipid metabolism. These data provided a comprehensive picture of genetic factors mediated molecular differences in the immune response to influenza infection between sexes.

Sex-differentiated gene expression in normal and disease states of human tissues has been identified in many studies [52–56]. However, the interpretation of underlying molecular mechanisms of infectious disease pathogenesis, especially genetic factors mediated baseline differences between the sexes, remains challenging due to the inevitable presence of confounding factors such as sex hormones, age, microbiota composition, and nutritional status in human studies. With the neutered adult ferrets, our study provided a unique opportunity to interrogate sex-biased transcriptional response to influenza virus infection driven by genetic mediators. By characterizing the blood transcriptome differences, we detected a temporal difference in the dynamics of immune responses between sexes, manifested by a more rapid activation in females than males upon infection. The immediate response in females is likely attributed to germline-encoded bias in innate immune sensing, as *TLR7* that is located on the X chromosome and encodes an innate PRR recognizing single-stranded RNA can not only have a higher expression level in females than males due to X-inactivation escape [57], but also induce more robust IFN-*α* production in peripheral blood mononuclear cells (PBMCs) isolated from women than those from men *in vitro* [58]. Moreover, we found limited overlap in the key regulators modulating commonly altered processes upon infection between sexes, suggesting differential regulation of various aspects of immune responses in females and males. This observation is consistent with a previous report of sex differences in regulatory targeting patterns identified across multiple healthy tissues in humans [54].

Using a network-based approach, we identified sex-specific modules that represented distinct pathways differentially impacted between sexes and exhibited unique regulatory structures and expression patterns. These modules revealed several sex-biased processes involved in the UPR, lipid metabolism, and inflammation. We found a robust early activation of IRE1/XBP1-dependent UPR in the female ferrets. Activation of IRE1 can lead to the splicing of XBP1 mRNA (XBP1u) and subsequent expression of spliced XBP1 (XBP1s) that acts as an active and stable transcription factor for UPR-related genes [39, 40]. Extensive studies on the role of UPR in immune cells demonstrated a fundamental impact of the IRE1/XBP1 pathway on promoting innate pro-inflammatory responses following TLR signaling in macrophages [59] and on modulating differentiation of several cell types (reviewed in [39]), including stimulating the differentiation of B cells into antibody-secreting plasma cells [60]. For example, the UPR is known to play a role in influenza virus infection [61]. Although predominantly observed in the lung, influenza virus infection induces ER-stress and activates the IRE1 pathway with subsequent XBP1 splicing [62]. Interestingly, emerging evidence from glycomic profiling of lung tissues collected from influenza-infected adult female ferrets indicates a link between the UPR, glycosylation, augmented immune response, and severe infection outcome; the activation of the IRE1/XBP1 pathway induces high mannose immediately upon infection, which can be recognized by the innate immune activator mannan-binding lectin (MBL2) and is associated with alveolar severity [63]. Moreover, a higher level of XBP1s expression in rat livers from prepubertal females compared with age-matched males has been reported, suggesting genetically determined female bias in XBP1s expression [64]. Taken together, rapid activation of the IRE1/XBP1 axis of the UPR observed specifically in the females, likely due to genetically encoded bias in XBP1s expression, may trigger potent immune and inflammatory responses and eventually could have contributed to a more severe disease seen in females than in males. Future studies will need to determine the functional consequences of XBP1s activation on a specific immune cell type during influenza infection and evaluate the impact of sex hormones on the pathways outlined.

Our integrative network analysis also captured two male-biased signaling pathways related to inflammatory response and lipid metabolism. Genes involved in the IL-1 pathway, including *IL1RAP, JUN, IRAK3*, and *IRAK4*, as well as two key regulators (*ACSL3* and *IL18RAP*), exhibited stronger inductions from 3 dpi onwards in males. ACSL3, primarily expressed in the endoplasmic reticulum (ER) and cytosolic lipid droplets, is responsible for fatty acid (FA) activation by converting free FA to FA acyl-CoA esters, an initial step contributing to lipid biosynthesis and FA oxidation [47, 65, 66]. Interestingly, lipid metabolism has been shown to dramatically alter the phenotype and function of monocyte/macrophages and modulate inflammatory responses, as increased lipid synthesis fueled by glucose metabolism has been linked to inflammatory macrophages, and FA oxidation is associated with alternatively activated macrophages [48, 49, 67]. However, the impact of ACSL3 induction and lipid metabolism on inflammatory responses in the blood cells and the crosstalk between the metabolic reprogramming and IL-1 family member IL-18-mediated signaling require further investigation. In addition to genes involved in the IL-1 pathway, we found another module enriched for the AP-1 pathway with a male-specific regulatory pattern. However, the extent of change in expression of its key regulators upon infection is similar between sexes. The AP-1 pathway comprised of members in the Jun, Fos, Maf, and ATF subfamilies [68, 69], plays an important role in regulating immune and inflammatory responses such as T-cell activation and cytokine production [70–73]. Intriguingly, two key regulators in this AP-1 pathway-related module — MS4A2 acting to amplify Fc*ε*RI expression and signaling during allergic responses [74] and HPGD responsible for inactivating potent lipid mediator PGE_2_ [45] — have been implicated in susceptibility to anaphylaxis by modulating PGE_2_ homeostasis in the mast cells (MCs) [44]. PGE_2_ stabilization by inhibiting HPGD-catalyzed degradation can attenuate Fc*ε*RI-mediated MC degranulation, preventing MC hyperresponsiveness and reducing susceptibility to anaphylaxis [44]. Thus, these observations likely indicate genetically encoded male bias in the expression and regulation pattern of genes involved in specific aspects of lipid metabolism and inflammatory response in the blood cells, which requires further investigation.

Taken together, our data revealed an emerging picture in which genetic factors mediated sex differences could manifest in the temporal dynamics and regulatory relationship of commonly induced immune and inflammatory responses, the activation of the IRE1/XBP1 pathway during the UPR, and the modulation of lipid metabolism and inflammatory response involving the IL-1 and AP-1 pathways. Although this proposed paradigm awaits further testing, especially the assessment of its generalizability in the presence of sex hormones, it provides insights into the molecular mechanism of sex differences in the blood cell function and activity, which can also be relevant to the pathogenesis of other infectious and inflammatory diseases. We noticed several limitations in this study, including the lack of granularity in differentiating cell type-specific responses and a restricted time course that does not allow a thorough investigation of adaptive immune responses, particularly humoral-mediated immunity. However, our study represents an important step towards a comprehensive understanding of sex differences in influenza virus-induced pathogenesis, which holds promise in developing personalized therapeutic strategies with improved efficacy for both sexes.

## MATERIALS AND METHODS

### Animal experiments

Female and male Fitch ferrets (*Mustela putorius furo*) were obtained from Triple F Farms (Sayre, PA, USA) and were seronegative to a circulating influenza A/California/07/2009 (A/CA/07/09; H1N1pdm09) virus. As previously described [75], adult ferrets, 6 to 12 months of age, were pair-housed in stainless steel cages (Shor-Line, Kansas City, KS) that contained Sani-Chips laboratory animal bedding (P.J. Murphy Forest Products, Montville, NJ) and provided with food and fresh water *ad libitum*. To achieve a mean of 20% weight loss no earlier than 8 dpi, adult ferrets were infected intranasally with the H1N1pdm09 virus at a dose of 10^6^ plaque-forming units (PFU). The animals were monitored daily for the severity of clinical disease using weight loss. Disease symptoms, including elevated temperature, low activity level, sneezing, and nasal discharge, were noted if present (data not shown). Any animal losing >20% weight loss was humanely euthanized. Ferrets were randomly assigned to be sacrificed at 1, 3, 5, 8, or 14 (only for females) dpi unless their clinical conditions (e.g., loss of >20% body weight) required a humane endpoint. Blood was collected from anesthetized ferrets via the anterior vena cava after infection. The University of Georgia Institutional Animal Care and Use Committee approved all experiments, which were conducted in accordance with the NIH’s Guide for the Care and Use of Laboratory Animals [75], The Animal Welfare Act, and the Biosafety in Microbiological and Biomedical Laboratories guide of the Centers for Disease Control and Prevention and the NIH.

### Ethics statement

All research studies involving the use of animals were reviewed and approved by the Institutional Animal Care and Use Committees of the University of Georgia. Studies were carried out in strict accordance with the recommendations in the Guide for the Care and Use of Laboratory Animals.

### Viral plaque assay

Plaque assays were performed to determine the viral burden in snap-frozen nasal washes, as previously described in [75]. Supernatants were gently thawed on ice, forced through a cell strainer (70 μm) and syringe plunger in phosphate-buffered saline, then spun down (2500 rpm, 5 minutes, 4°C) to collect the supernatant. Supernatants were diluted in Iscove’s modified Dulbecco’s minimum essential medium. Madin-Darby canine kidney (MDCK) cells were plated (5 × 10^5^) in each well of a six-well plate. Samples were diluted in Iscove’s modified Dulbecco’s minimum essential medium (final dilution factors of 10^0^ to 10^-6^) and overlaid onto the cells in 100 μL of Dulbecco’s modified Eagle’s medium supplemented with penicillin-streptomycin followed by 1 hour incubation. Samples were removed, cells were washed twice, and medium was replaced with 2 ml of L15 medium plus 0.8% agarose (Cambrex, East Rutherford, NJ, USA) and incubated for 72 hours at 37°C with 5% carbon dioxide. Agarose was removed and discarded. The cells were fixed with 10% buffered formalin and then stained with 1% crystal violet for 15 minutes. After thorough washing in distilled water to remove excess crystal violet, the plates were dried, the number of plaques was counted, and the number of PFU per milliliter was calculated.

### RNA-seq experiments

RNA was extracted from ferret blood using the Mouse RiboPure-Blood RNA Isolation Kit (Ambion). Both extraction methods followed the manufacturer’s protocols and incorporated a DNase treatment (QIAGEN) after passing the sample through the filter cartridge. Strand-specific total RNA-seq libraries from ribosomal RNA-depleted RNA were prepared using the TruSeq Stranded Total RNA Library Prep kit (Illumina) according to the manufacturer-supplied protocol. Libraries were sequenced 100 bp paired-end to a depth of approximately 40 million host genome reads on Illumina HiSeq 2500 instruments.

### RNA-seq analysis

Paired reads were aligned to the ferret genome Ensembl version 1.0.80 using tophat (version v2.0.13). Following read alignment, featureCounts (v1.4.6) was used to quantify the mRNA expression levels based on the Ensembl gene model. mRNAs with more than 5 reads in at least 1 sample were considered expressed and retained for further analysis. Otherwise, they were removed. The mRNA read counts data was normalized using the trimmed mean of M-values normalization (TMM) method of the edgeR package [76] to adjust for sequencing library size difference.

### Identification of differentially expressed genes

Given the time series information for each different dataset, we used two distinct methods to identify differentially expressed genes. We used a one-way ANOVA-like approach as implemented in the Limma-package [77] to identify differentially expressed genes across the time series—termed significant response genes. Significance is defined based on a false discovery rate (FDR) of 5% or less. DEGs is used as a generic term for differentially expressed genes, including SRGs. Statistical significance of intersections between any two given DEG sets in both sexes was determined with SuperExactTest [78].

### Gene co-expression network analysis

Multiscale Embedded Gene Co-Expression Network Analysis (MEGENA) [35] was performed to identify host modules of highly co-expressed genes in influenza infection. The MEGENA workflow comprises four major steps: 1) Fast Planar Filtered Network construction (FPFNC), 2) Multiscale Clustering Analysis (MCA), 3) Multiscale Hub Analysis (MHA), 4) and Cluster-Trait Association Analysis (CTA). The total relevance of each module to influenza virus infection was calculated by using the Product of Rank method with the combined enrichment of the differentially expressed gene (DEG) signatures as implemented: G_j_ = Π_i_ g_ji_, where, g_ji_ is the relevance of a consensus **j** to a signature **i;** and g_ji_ is defined as (max_j_(r_ji_) + 1 − r_ji_)/∑_j_ r_ji_, where r_ji_ is the ranking order of the significance level of the overlap between module **j** and the signature.

### Identification of enriched pathways and key regulators in the host transcriptome modules

To functionally annotate gene signatures and gene modules derived from the host transcriptome data, we performed an enrichment analysis of the established pathways and signatures—including the MSigDB and ARCHS^4^ tissues [79]. The hub genes in each subnetwork were identified using the adopted Fisher’s inverse Chi-square approach in MEGENA; Bonferroni-corrected p-values smaller than 0.05 were set as the threshold to identify significant hubs.

### Correlation analysis

The correlation between modules and physiological traits was performed using Spearman’s correlation. The correlation between the FCs of DEGs in females and males over time was estimated using Pearson’s correlation with the Benjamini-Hochberg correction.

### Similarity comparison of modules and their key drivers

The pairwise similarity between a given female module and every male module or between their key drivers was estimated using Fisher’s Exact Test (FET) with Bonferroni correction. Modules in one sex without counterparts in the other sex (FET p ≤ 0.05) were considered sex-unique module compositions. Differential Gene Correlation Analysis (DGCA) was also performed with the “moduleDC” function implemented in the DGCA package [38] to further determine the unique regulatory relationship of the overlapped gene members between a pair of similar modules in females and males. Modules with significant differential correlation (p ≤ 0.05) with all of their counterparts were considered to exhibit sex-specific regulatory relationships.

## CODE AVAILABILITY

Computer code, scripts, and provenance information to produce the data discussed in this manuscript are available at Synapse as part of the FluDyNeMo project (https://www.synapse.org/#!Synapse:syn25049511).

## DATA AVAILABILITY

RNAseq data have been submitted to GEO/SRA and are accessible under accession numbers GSE168512. Ferret transcriptome information and derived data are also available at Synapse as part of the FluDyNeMo project (http://www.synapse.org/fludynemo).

## AUTHOR CONTRIBUTIONS

C.V.F. conceived the project, C.W. and C.V.F. performed the computational study, M.W. helped with RNAseq data processing, E.G., and T.M.R., designed the experiments, L.P.L., C.E.C. and S.K.J. executed the experiments, B.Z. helped with the computational analysis and provided guidance, C.W. and C.V.F. wrote the manuscript. All the authors reviewed the manuscript.

## Supplementary Figures

**Figure S1.**
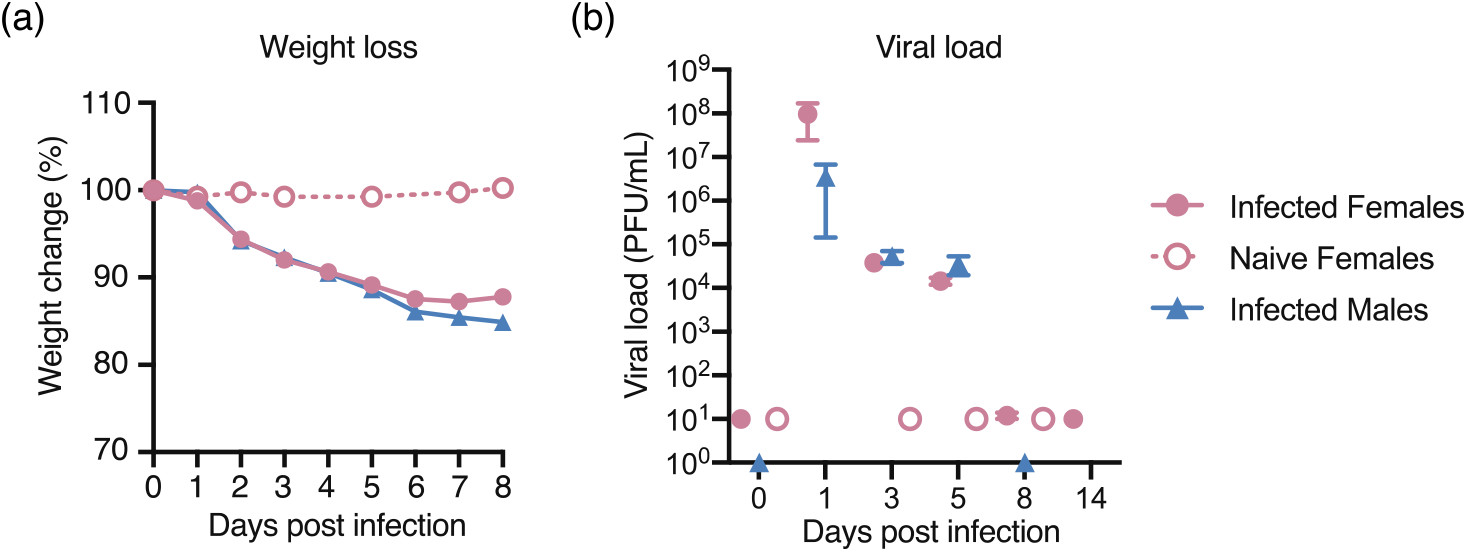
Physiological traits revealing pathogenicity of influenza infection in both sexes. (**a**) Weight change was monitored daily for 8 days post-infection (n = 4 for naïve females, n = 80 for infected females, and n = 32 for infected males). (**b**) The viral titers in the nasal washes were determined by plaque assays with MDCK cells (n = 4 for naïve females, n = 49 for infected females, and n = 32 for infected males). The dots represent the mean, and the error bars indicate the standard error of the mean (SEM).

**Figure S2.**
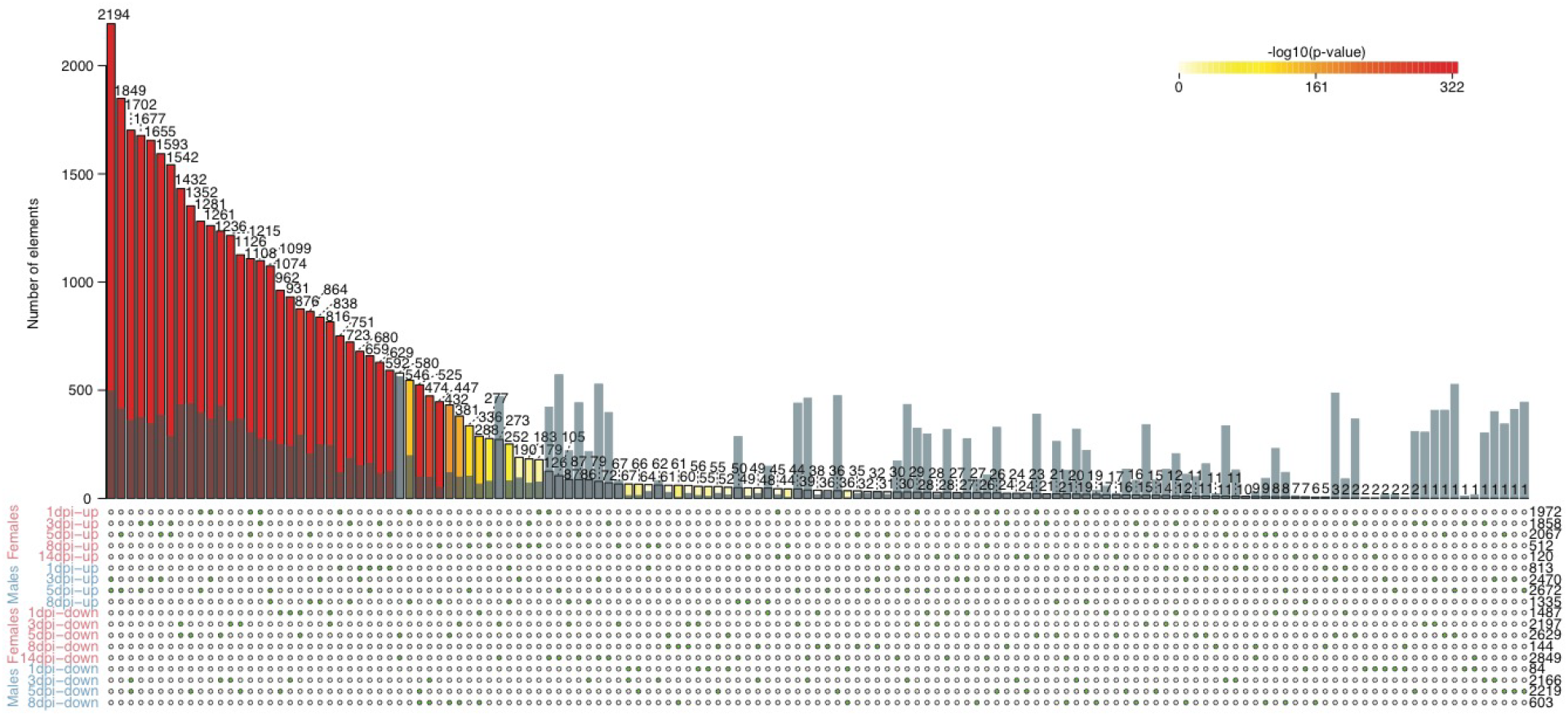
Pairwise comparison of up- or down-regulated differentially expressed genes (DEGs) at each time point in both sexes. DEGs were obtained by comparing the infected samples with their controls for each sex. The solid bars are colored by -log10(p-value) generated with the super exact test and the numbers on top show the observed intersects of DEGs between a pair of DEG sets. The semi-transparent blue bars overlaying the solid bars indicate the expected intersects. The green dots below denote two sets of DEGs used in the comparisons. The numbers to the right of the dot matrix indicate the number of DEGs in a given set.

**Figure S3.**
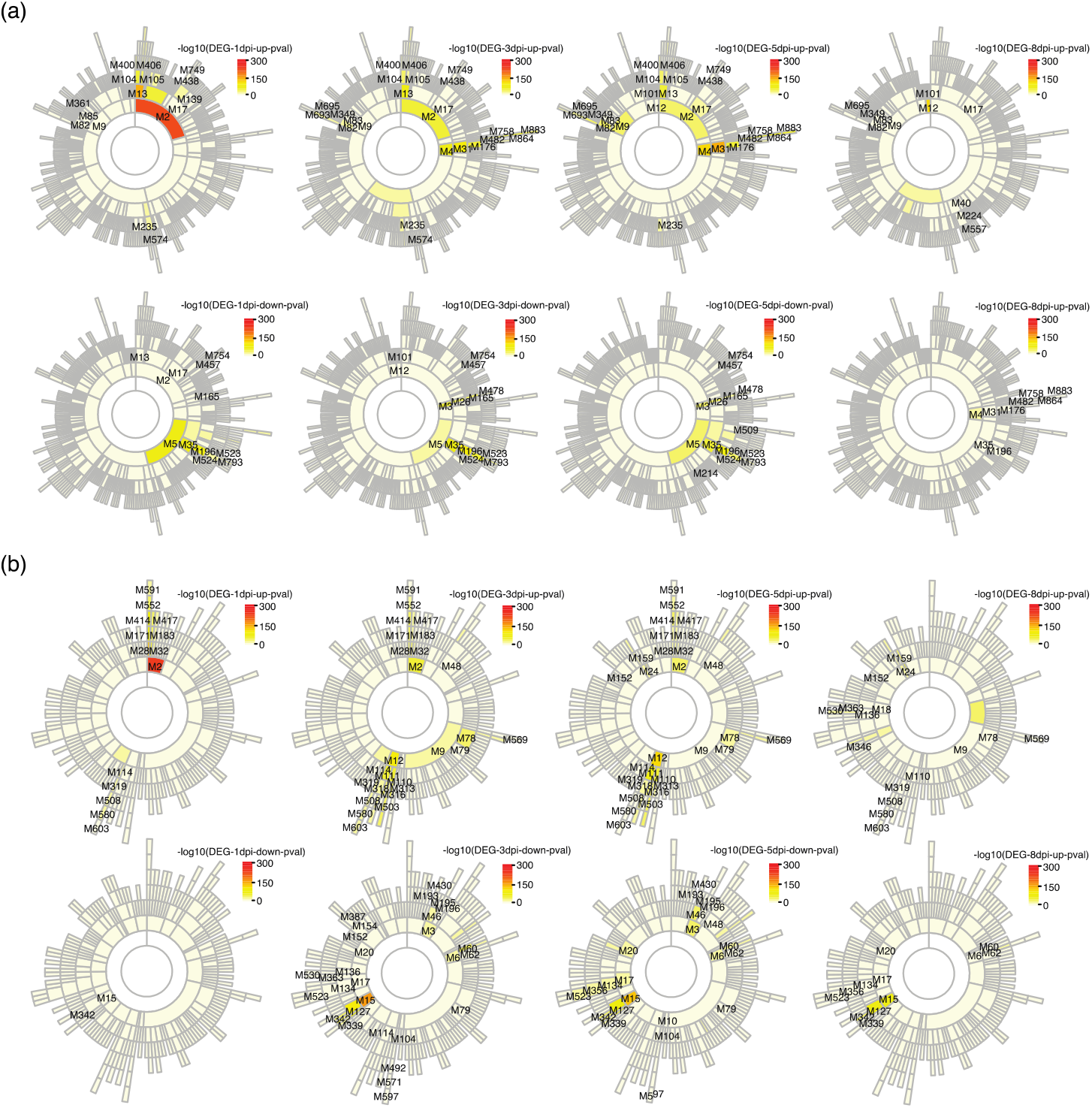
Sunburst plots showing the module hierarchy in the (a) female and (b) male co-expression networks and highlighting those enriched in differentially expressed genes (DEGs) and the Molecular Signatures Database (MSigDB) gene sets. The colors indicate the -log10(p-value) for Fisher’s exact test of enrichment of up- or down-regulated DEGs at each time point.

**Figure S4.**
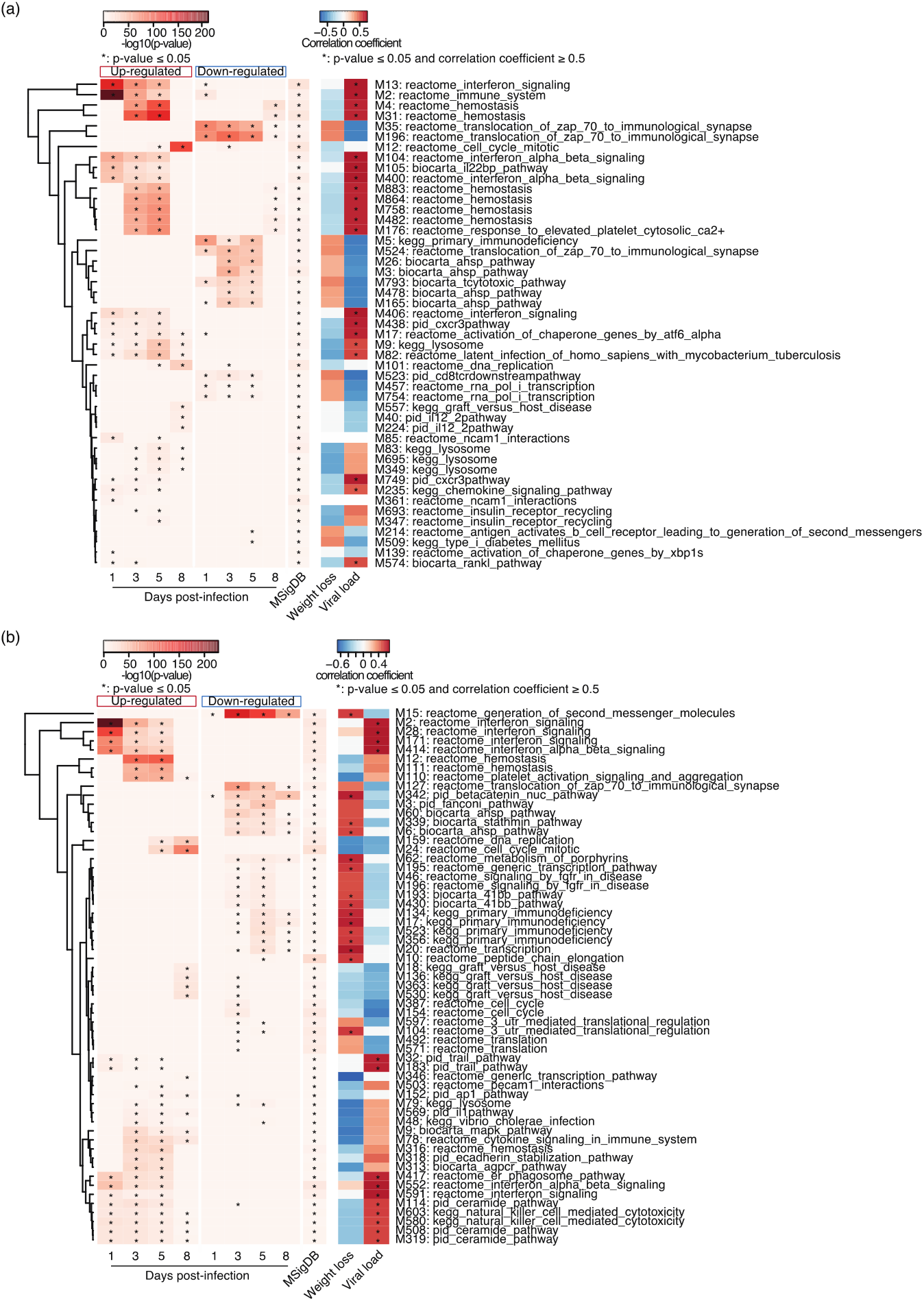
Heatmaps showing the enrichment of DEGs in modules and module-trait correlation in the (a) female and (b) male networks. The first three panels on the left side of the plot show the -log10(p-value) obtained from fisher’s exact test of enrichment of up- or down-regulated DEGs or the Molecular Signatures Database (MSigDB) gene sets at each time point for modules indicated on the right side. The asterisks in these panels denote p ≤ 0.05. The fourth panel shows Spearman’s correlation coefficient between modules and traits. The asterisks in this panel denote p ≤ 0.05 and correlation coefficient ≥ 0.5.

**Figure S5.**
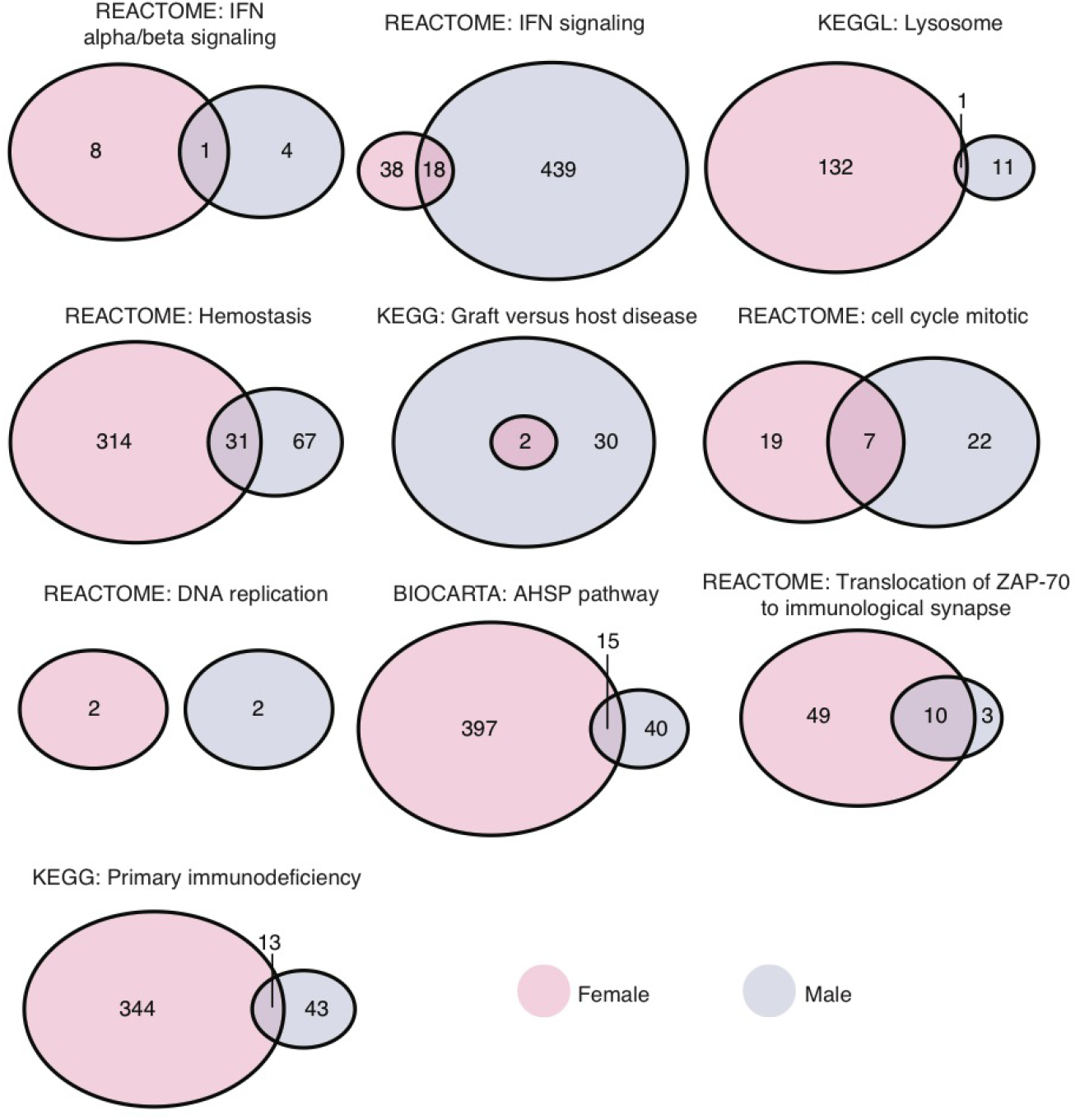
Comparison of key regulators in modules enriched in the Molecular Signatures Database (MSigDB) gene sets commonly observed in both sexes. All the key regulators in modules enriched in a given MSigDB gene set in each sex were used for the comparison. The color of circles indicates the sex (pink for females and blue for males).

**Figure S6.**
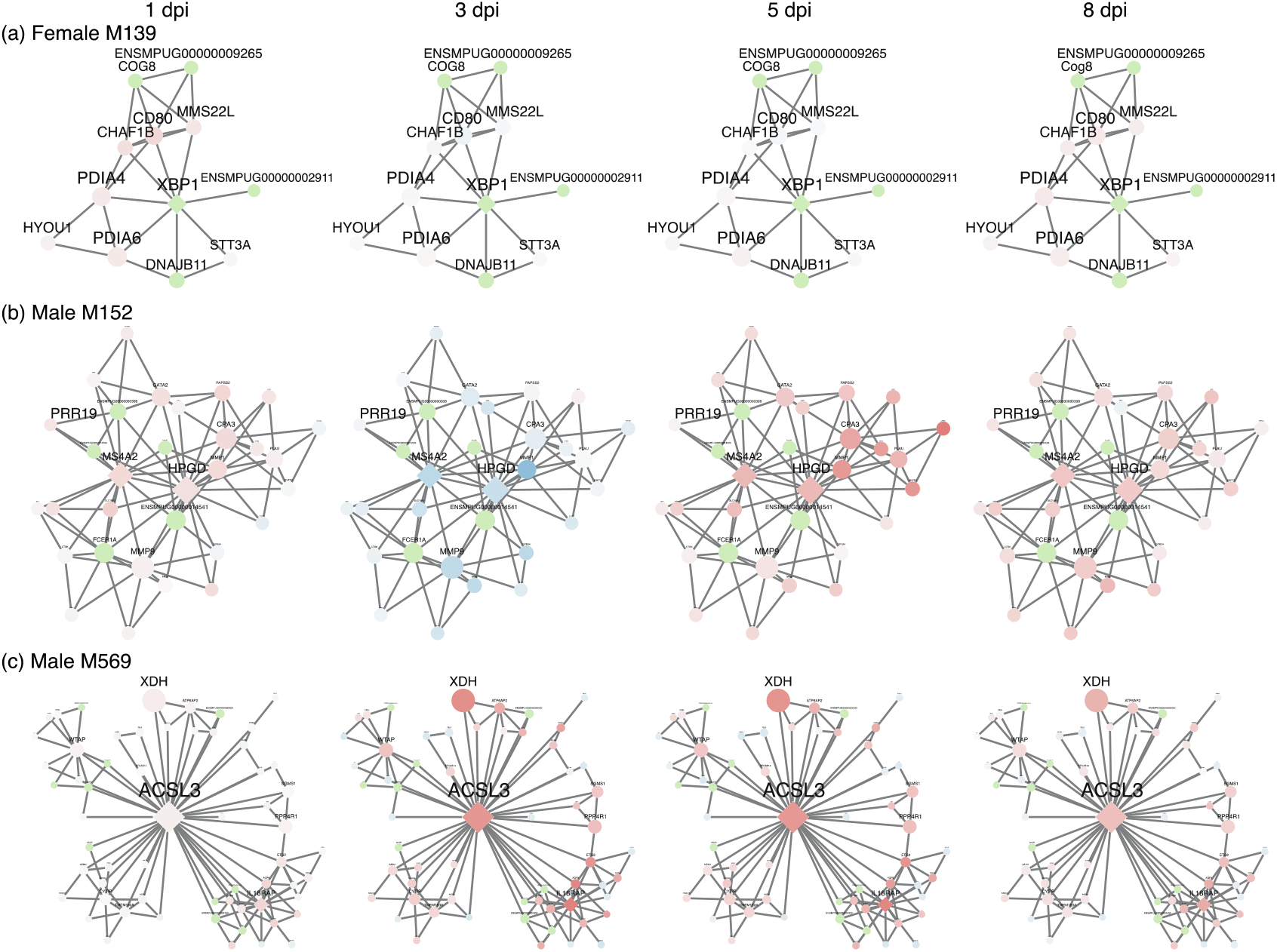
Differentially expressed genes in sex-specific modules. Networks of (**a**) the female-specific module M139 and male-specific modules (**b**) M152 and (**c**) M569. Nodes were colored by their fold change over time (red for up-regulation, blue for down-regulation, and light green for no significant differential expression). Node shapes denote the module membership (diamonds for key regulators and circles for regular members), and their sizes are proportional to the node strength.

## REFERENCES

1. Klein, S.L. and K.L. Flanagan, Sex differences in immune responses. Nature Reviews Immunology, 2016. 16(10): p. 626–38.

2. Jacobson, D.L., et al., Epidemiology and estimated population burden of selected autoimmune diseases in the United States. Clinical Immunology and Immunopathology, 1997. 84(3): p. 223–243.

3. Cook, M.B., et al., Sex disparities in cancer incidence by period and age. Cancer Epidemiology, Biomarkers & Prevention, 2009. 18(4): p. 1174–1182.

4. Cook, M.B., et al., Sex disparities in cancer mortality and survival. Cancer Epidemiology, Biomarkers & Prevention, 2011. 20(8): p. 1629–1637.

5. Kim, H.-I., H. Lim, and A. Moon, Sex Differences in Cancer: Epidemiology, Genetics and Therapy. Biomolecules & Therapeutics, 2018. 26(4): p. 335–342.

6. vom Steeg, L.G. and S.L. Klein, SeXX matters in infectious disease pathogenesis. PLoS Pathogens, 2016. 12(2): p. e1005374.

7. Fischer, J., et al., Sex differences in immune responses to infectious diseases. Infection, 2015. 43(4): p. 399–403.

8. Sawyer, C.C., Child mortality estimation: Estimating sex differences in childhood mortality since the 1970s. PLoS Medicine, 2012. 9(8): p. e1001287.

9. Giefing-Krôll, C., et al., How sex and age affect immune responses, susceptibility to infections, and response to vaccination. Aging Cell, 2015. 14: p. 309–321.

10. Klein, S.L., Sex influences immune responses to viruses, and efficacy of prophylaxis and treatments for viral diseases. Bioessays, 2012. 34(12): p. 1050–9.

11. Morgan, R. and S.L. Klein, The intersection of sex and gender in the treatment of influenza. Current Opinion in Virology, 2019. 35: p. 35–41.

12. WHO, Sex, gender and influenza. 2010. p. 58.

13. Lorenzo, M.E., et al., Antibody responses and cross protection against lethal influenza A viruses differ between the sexes in C57BL/6 mice. Vaccine, 2011. 29(49): p. 9246–9255.

14. Fink, A.L., et al., Biological sex affects vaccine efficacy and protection against influenza in mice. PNAS, 2018. 115(49): p. 12477–12482.

15. Potluri, T., et al., Age-associated changes in the impact of sex steroids on influenza vaccine responses in males and females. NPJ Vaccines, 2019. 4: p. 29.

16. Scully, E.P., et al., Considering how biological sex impacts immune responses and COVID-19 outcomes. Nature Reviews Immunology, 2020. 20(7): p. 442–447.

17. Takahashi, T., et al., Sex differences in immune responses that underlie COVID-19 disease outcomes. Nature, 2020. 588(7837): p. 315–320.

18. Galligan, C.L. and E.N. Fish, Sex Differences in the Immune Response, in Sex and Gender Differences in Infection and Treatments for Infectious Diseases, S.L. Klein and C.W. Roberts, Editors. 2015, Springer, Cham. p. 1–29.

19. Robinson, D.P., et al., Elevated 17beta-estradiol protects females from influenza A virus pathogenesis by suppressing inflammatory responses. PLoS Pathogens, 2011. 7(7): p. e1002149.

20. Markle, J.G.M., et al., Sex differences in the gut microbiome drive hormone-dependent regulation of autoimmunity. Science, 2013. 339(6123): p. 1084–1088.

21. Yurkovetskiy, L., et al., Gender bias in autoimmunity is influenced by microbiota. Immunity, 2014. 39(2): p. 400–412.

22. Vom Steeg, L.G. and S.L. Klein, Sex steroids mediate bidirectional interactions between hosts and microbes. Hormones and Behavior, 2017. 88: p. 45–51.

23. Vemuri, R., et al., The microgenderome revealed: Sex differences in bidirectional interactions between the microbiota, hormones, immunity and disease susceptibility. Seminars in Immunopathology, 2019. 41(2): p. 265–275.

24. Khulan, B., et al., Periconceptional maternal micronutrient supplementation is associated with widespread gender related changes in the epigenome: a study of a unique resource in the Gambia. Human Molecular Genetics, 2012. 21(9): p. 2086–2101.

25. Tobi, E.W., et al., DNA methylation differences after exposure to prenatal famine are common and timing- and sex-specific. Human Molecular Genetics, 2009. 18(21): p. 4046–4053.

26. Sinha, A., et al., Reduced risk of neonatal respiratory infections among breastfed girls but not boys. Pediatrics, 2003. 112(4): p. e303.

27. Kawai, K., et al., Sex differences in the effects of maternal vitamin supplements on mortality and morbidity among children born to HIV-infected women in Tanzania. British Journal of Nutrition, 2010. 103(12): p. 1784–1791.

28. Osrin, D., et al., Effects of antenatal multiple micronutrient supplementation on birthweight and gestational duration in Nepal: double-blind, randomised controlled trial. Lancet, 2005. 365(9463): p. 955–962.

29. Christoforidou, Z., et al., Sexual dimorphism in immune development and in response to nutritional intervention in neonatal piglets. Frontiers in Immunology, 2019. 10: p. 2705.

30. Fish, E.N., The X-files in immunity: Sex-based differences predispose immune responses. Nature Reviews Immunology, 2008. 8: p. 737–744.

31. Klein, S.L., The effects of hormones on sex differences in infection: From genes to behavior. Neuroscience and Biobehavioral Reviews, 2000. 24: p. 627–638.

32. Schurz, H., et al., The X chromosome and sex-specific effects in infectious disease susceptibility. Human Genomics, 2019. 13(1): p. 2.

33. Bianchi, I., et al., The X chromosome and immune associated genes. Journal of Autoimmunity, 2012. 38: p. J187–J192.

34. Rowe, T., et al., Modeling host responses in ferrets during A/California/07/2009 influenza infection. Virology, 2010. 401(2): p. 257–65.

35. Song, W.M. and B. Zhang, Multiscale Embedded Gene Co-expression Network Analysis. PLoS Comput Biol, 2015. 11(11): p. e1004574.

36. Le, V.B., et al., Platelet activation and aggregation promote lung inflammation and influenza virus pathogenesis. American Journal of Respiratory and Critical Care Medicine, 2015. 191(7): p. 804–19.

37. Ren, C., et al., Identification and characterization of RTVP1/GLIPR1-like genes, a novel p53 target gene cluster. Genomics, 2006. 88(2): p. 163–72.

38. McKenzie, A.T., et al., DGCA: A comprehensive R package for Differential Gene Correlation Analysis. BMC Systems Biology, 2016. 10: p. 106.

39. So, J.S., Roles of endoplasmic reticulum stress in immune responses. Molecules and Cells, 2018. 41(8): p. 705–716.

40. Hetz, C., The unfolded protein response: controlling cell fate decisions under ER stress and beyond. Nature Reviews Molecular Cell Biology, 2012. 13(2): p. 89–102.

41. Ugajin, T., et al., Fc epsilonRI, but not FcgammaR, signals induce prostaglandin D2 and E2 production from basophils. The American Journal of Pathology, 2011. 179(2): p. 775–82.

42. Inage, E., et al., Critical Roles for PU. 1, GATA1, and GATA2 in the expression of human FcepsilonRI on mast cells: PU. 1 and GATA1 transactivate FCER1A, and GATA2 transactivates FCER1A and MS4A2. The Journal of Immunology, 2014. 192(8): p. 3936–46.

43. Mkaddem, S.B., M. Benhamou, and R.C. Monteiro, Understanding Fc receptor involvement in inflammatory diseases: From mechanisms to new therapeutic tools. Frontiers in Immunology, 2019. 10: p. 811.

44. Rastogi, S., et al., PGE_2_ deficiency predisposes to anaphylaxis by causing mast cell hyperresponsiveness. The Journal of Allergy and Clinical Immunology, 2020. 146(6): p. 1387–1396.

45. Tai, H.-H., et al., NAD+-linked 15-hydroxyprostaglandin dehydrogenase: Structure and biological functions. Current Pharmaceutical Design, 2006. 12(8): p. 955–962.

46. Yan, S., et al., Long-chain acyl-CoA synthetase in fatty acid metabolism involved in liver and other diseases: an update. World Journal of Gastroenterology, 2015. 21(12): p. 3492–8.

47. Roelands, J., et al., Long-chain acyl-CoA synthetase 1 role in sepsis and immunity: Perspectives from a parallel review of public transcriptome datasets and of the literature. Frontiers in Immunology, 2019. 10: p. 2410.

48. Remmerie, A. and C.L. Scott, Macrophages and lipid metabolism. Cellular Immunology, 2018. 330: p. 27–42.

49. Batista-Gonzalez, A., et al., New insights on the role of lipid metabolism in the metabolic reprogramming of macrophages. Frontiers in Immunology, 2020. 10: p. 2993.

50. Kobori, T., et al., Interleukin-18 amplifies macrophage polarization and morphological alteration, leading to excessive angiogenesis. Frontiers in Immunology, 2018. 9: p. 334.

51. Yasuda, K., K. Nakanishi, and H. Tsutsui, Interleukin-18 in health and disease. International Journal of Molecular Sciences, 2019. 20(3): p. 649.

52. Kassam, I., et al., Tissue-specific sex differences in human gene expression. Human Molecular Genetics, 2019. 28(17): p. 2976–2986.

53. Trabzuni, D., et al., Widespread sex differences in gene expression and splicing in the adult human brain. Nature Communications, 2013. 4: p. 2771.

54. Lopes-Ramos, C.M., et al., Sex differences in gene expression and regulatory networks across 29 human tissues. Cell Reports, 2020. 31(12): p. 107795.

55. Bongen, E., et al., Sex Differences in the Blood Transcriptome Identify Robust Changes in Immune Cell Proportions with Aging and Influenza Infection. Cell Reports, 2019. 29(7): p. 1961–1973.

56. Jansen, R., et al., Sex differences in the human peripheral blood transcriptome. BMC Genomics, 2014. 15: p. 33.

57. Pisitkun, P., et al., Autoreactive B cell responses to RNA-related antigens due to TLR7 gene duplication. Science, 2006. 312(5780): p. 1669–1672.

58. Berghöher, B., et al., TLR7 ligands induce higher IFN-alpha production in females. The Journal of Immunology, 2006. 177(4): p. 2088–2096.

59. Martinon, F., et al., TLR activation of the transcription factor XBP1 regulates innate immune responses in macrophages. Nature Immunology, 2010. 11: p. 411–418.

60. Iwakoshi, N.N., et al., Plasma cell differentiation and the unfolded protein response intersect at the transcription factor XBP-1. Nature Immunology, 2003. 4(4): p. 321–329.

61. Mehrbod, P., et al., The roles of apoptosis, autophagy and unfolded protein response in arbovirus, influenza virus, and HIV infections. Virulence, 2019. 10(1): p. 376–413.

62. Hassan, I.H., et al., Influenza A viral replication is blocked by inhibition of the inositol-requiring enzyme 1 (IRE1) stress pathway. Journal of Biological Chemistry, 2012. 287(7): p. 4679–4689.

63. Heindel, D.W., et al., Glycomic analysis of host response reveals high mannose as a key mediator of influenza severity. PNAS, 2020. 117(43): p. 26926–26935.

64. Rossetti, C.L., et al., Sexual dimorphism of liver endoplasmic reticulum stress susceptibility in prepubertal rats and the effect of sex steroid supplementation. Experimental Physiology, 2019. 104: p. 677–690.

65. Yan, S., et al., Long-chain acyl-CoA synthetase in fatty acid metabolism involved in liver and other diseases: An update. World Journal of Gastroenterology, 2015. 21(12): p. 3492–3498.

66. Tang, Y., et al., Fatty acid activation in carcinogenesis and cancer development: Essential roles of long-chain acyl-CoA synthetases. Oncology Letters, 2018. 16(2): p. 1390–1396.

67. Yan, J. and T. Horng, Lipid metabolism in regulation of macrophage functions. Trends in Cell Biology, 2020. 30(12): p. 979–989.

68. Eferl, R. and E.F. Wagner, AP-1: a double-edged sword in tumorigenesis. Nature Reviews Cancer, 2003. 3: p. 859–868.

69. Shaulian, E. and M. Karin, AP-1 as a regulator of cell life and death. Nature Cell Biology, 2002. 4: p. E131–E136.

70. Atsaves, V., et al., AP-1 transcription factors as regulators of immune responses in cancer. Cancers (Basel), 2019. 11(7).

71. Zenz, R., et al., Activator protein 1 (Fos/Jun) functions in inflammatory bone and skin disease. Arthritis Research & Therapy, 2008. 10(1): p. 201.

72. Foletta, V.C., D.H. Segal, and D.R. Cohen, Transcriptional regulation in the immune system: all roads lead to AP-1. Journal of Leukocyte Biology, 1998. 63: p. 139–152.

73. Schonthaler, H.B., J. Guinea-Viniegra, and E.F. Wagner, Targeting inflammation by modulating the Jun/AP-1 pathway. Annals of the Rheumatic Diseases, 2011. 70: p. i109–i112.

74. Kraft, S., et al., The role of the Fc*ε*RI *β*-chain in allergic diseases. International Archives of Allergy and Immunology, 2004. 135: p. 62–72.

75. Bissel, S.J., et al., Age-Related Pathology Associated with H1N1 A/California/07/2009 Influenza Virus Infection. Am J Pathol, 2019.

76. Robinson, M.D., D.J. McCarthy, and G.K. Smyth, edgeR: a Bioconductor package for differential expression analysis of digital gene expression data. Bioinformatics, 2010. 26(1): p. 139–40.

77. Smyth, G.K., Linear models and empirical bayes methods for assessing differential expression in microarray experiments. Stat Appl Genet Mol Biol, 2004. 3: p. Article3.

78. Wang, M., Y. Zhao, and B. Zhang, Efficient test and visualization of multi-set intersections. Scientific Reports, 2015. 5: p. 16923.

79. Lachmann, A., et al., Massive mining of publicly available RNA-seq data from human and mouse. Nature communications, 2018. 9(1): p. 1–10.

